# Expression of recombinant human glutamylating TTLLs in human cells leads to differential tubulin glutamylation patterns, with only TTLL6 disrupting microtubule dynamics

**DOI:** 10.1101/2024.11.25.624814

**Authors:** Mohamed Aghyad Al Kabbani, Pragya Jatoo, Kathrin Klebl, Bert M. Klebl, Hans Zempel

**Author notes:** Correspondence to: Dr. Dr. Hans Zempel. Correspondence to: Dr. Pragya Jatoo.

## Abstract

Polyglutamylation, a post-translational modification (PTM) catalyzed by a subset of Tubulin Tyrosine Ligase-Like (TTLL) family enzymes, regulates microtubule dynamics through its influence on interactions with microtubule-associated, motor, and severing proteins, and has recently also been implicated in genetic and neurodegenerative diseases. In this study, we characterized the glutamylation activity of various glutamylating TTLLs in human embryonic kidney 293T (HEK293T) cells, revealing distinct patterns of mono- and polyglutamylation among TTLL family members, with TTLL4 and TTLL11 exhibiting the strongest chain initiation and elongation activities, respectively. We found that TTLL6 expression uniquely decreased microtubule stability, with live-cell imaging of end-binding protein (EB3) showing a TTLL6-induced decrease in microtubule stability. To explore therapeutic modulation of TTLL activity, we tested LDC10, a novel TTLL inhibitor, which successfully blocked glutamylation across all TTLLs investigated in this study, while also reversing the microtubule-destabilizing effects of TTLL6. These findings identify a potential pathogenic role of TTLL6 in microtubule dynamics and highlight LDC10 as a promising pharmacological tool to counteract TTLL-induced microtubule destabilization.

## Introduction

Most members of the Tubulin Tyrosine Ligase-Like (TTLL) family are responsible for glutamylation, a post-translational modification (PTM) that adds a glutamate side chain to a glutamate residue within the C-terminal tail of α- and β-tubulin (Janke et al., 2005). Some TTLLs, such as TTLL4 and TTLL7, initiate the side chain by adding the first glutamate, while others, including TTLL1, TTLL6 and TTLL11, elongate this side chain by adding additional residues (van Dijk et al., 2007).

Polyglutamylation is an important microtubule PTM as it regulates microtubule binding to microtubule-associated proteins (MAPs), motor, and severing (Genova et al., 2023). Long glutamate chains are known to recruit spastin and katanin, two microtubule severing enzymes, leading to microtubule severing (Lacroix et al., 2010). However, exceedingly long chains, above 8-9 glutamate residues, alter spastin’s function from severing to stabilizing the microtubules (Valenstein & Roll-Mecak, 2016).

Disruption of tubulin polyglutamylation homeostasis has been implicated in several diseases. Notably, hyperglutamylation resulting from deficient cytosolic carboxypeptidase 1 (CCP1), a major deglutamylase responsible for removing glutamate residues from polyglutamate side chains (Rogowski et al., 2010), is associated with neurodegeneration in mice and humans (Magiera et al., 2018; Shashi et al., 2018). Other disorders linked to TTLL dysfunction include impaired ciliary structure and function, retinal dystrophy, and male infertility (Bedoni et al., 2016; Kolawole et al., 2023; Oh et al., 2022; Vogel et al., 2010). However, therapeutic approaches targeting TTLLs are still lacking.

Here, we expressed different recombinant human glutamylating TTLLs in human embryonic kidney 293T (HEK293T) cells and observed distinct patterns of mono- and polyglutamylation. TTLL6 expression also disrupted microtubule stability, as shown through live-cell imaging of microtubule plus-end tracking protein EB3 comets. Interestingly, a novel TTLL inhibitor, LDC10, successfully blocked the glutamylating activity of all TTLLs investigated in this study and mitigated the destabilizing effect of TTLL6 on microtubules. In summary, we have identified a potential pathological role of TTLL6-mediated polyglutamylation and described a new pharmacological intervention to counteract it.

## Methods

### HEK293T cell maintenance, transfection, and inhibitor treatment

HEK293T cells were cultured in high glucose DMEM (Thermofisher Scientific) supplemented with 10% FBS and 1x Antibiotic/Antimycotic solution (Thermofisher Scientific) at 37 °C in a humidified incubator with 5% CO_2_. For EYFP-tagged TTLL expression, cells were seeded into 6-well plates and transfected with 3 µg DNA for 48 hours. For pharmacological treatments, cells were treated with 10 µM LDC10 or a vehicle control for 24 hours, beginning one day post-transfection.

### Western Blot

For Western blot analysis, HEK293T cells were lysed in RIPA buffer containing 1x protease & phosphatase inhibitor cocktail (Thermofisher Scientific). Lysates were diluted in 5x Laemmli sample buffer, boiled at 95°C for 5 minutes, and separated on 10% SDS-polyacrylamide gels. Proteins were then transferred to PVDF membranes, which were blocked for one hour in TBS-T containing 5% milk. Following blocking, membranes were incubated with the primary antibody overnight at 4°C, washed three times with TBS-T, and incubated with the corresponding HRP-conjugated secondary antibody for one hour at room temperature. After three additional TBS-T washes, immunoreactions were detected using SuperSignal West Femto Chemiluminescent Substrate (Thermofisher Scientific) and a ChemiDoc XRS + system (Bio-Rad).

### Live-cell imaging

For live-cell imaging, HEK293T cells were seeded into coated 6-well plates and co-transfected with 0.5 µg EB3-tdTomato and 1.5 µg EYFP-TTLL or empty EYFP vector. One day post-transfection, cells were transferred to a live-cell imaging chamber (ALA Scientific), and EB3 comets in single cells were imaged for 60 seconds (1 frame per 2 seconds) with a Leica DMi8 microscope (Leica). Only cells exhibiting both tdTomato and EYFP signals were included in the analysis. EB3 comet tracks were analyzed via ImageJ software using TrackMate plugin (Tinevez et al., 2017) as described in Allroggen et al., 2024. Microtubule dynamics were analyzed using LoG detector with an estimated object diameter of 1.5 µm, assessing parameters such as microtubule stability (s), microtubule run length (μm), and microtubule growth rate (μm/s).

### Antibodies

The antibodies used in this study are listed in Table. 1

**Table 1.** List of the antibodies used in this study.

| Antibody | Species | Clonality | Cat# | Supplier | RRID | Dilution |
| --- | --- | --- | --- | --- | --- | --- |
| Anti-Tubulin<br>Polyglutamylated<br>antibody (B3) | Mouse | Monoclonal | T9822 | Sigma-Aldrich | AB_477598 | 1:500 |
| Anti-<br>polyglutamylation<br>modification<br>(GT335) | Mouse | Monoclonal | AG-20B-0020 | AdipoGen | AB_2490211 | 1:500 |
| Anti-GFP<br>antibody | Rabbit | Polyclonal | ab290 | Abcam | AB_2313768 | 1:1000 |
| Anti-GAPDH<br>antibody (G-9) | Mouse | Monoclonal | sc-365062 | Santa Cruz | AB_10847862 | 1:1000 |
| Anti-beta actin<br>antibody | Mouse | Monoclonal | HRP-60008 | Proteintech | AB_2819183 | 1:5000 |

## Results

### Expression of different TTLLs reveals distinct patterns of tubulin glutamylation

Differential analysis of individual TTLLs is challenging, due to low endogenous expression and potentially overlapping glutamylation activity. Hence, to investigate the effects of different glutamylating TTLLs on tubulin glutamylation and microtubule dynamics, we expressed EYFP-tagged constructs of TTLL1, TTLL4, TTLL6, TTLL7, and TTLL11 in HEK293T cells for 2 days and assessed their expression levels via Western blotting. All recombinant TTLLs exhibited very low expression levels compared to the EYFP control (despite using the same transfection protocol including DNA amount), with EYFP-TTLL11 showing the highest expression (Fig. 1A).

**Figure 1.**
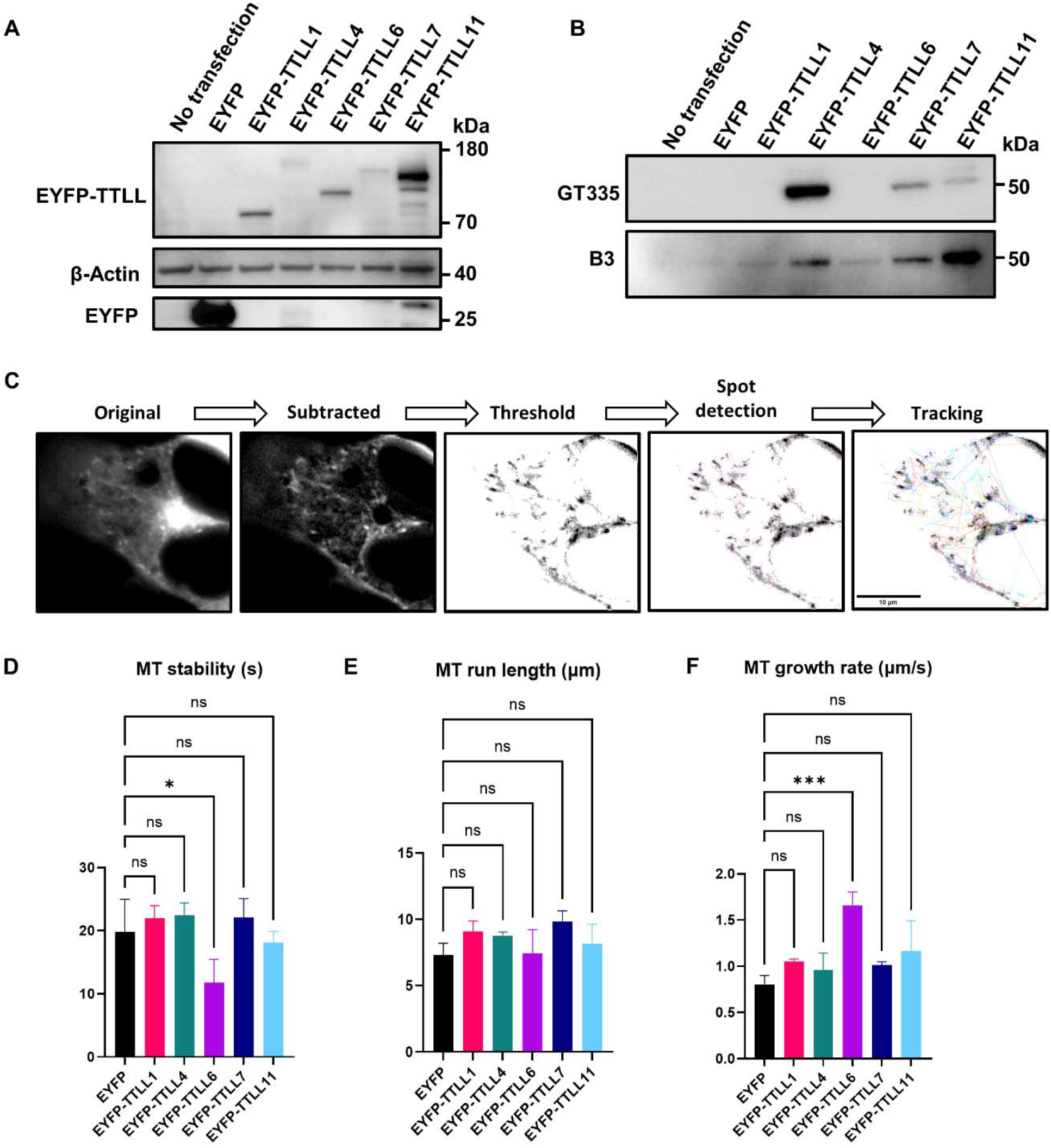
Expression of different TTLLs induces distinct glutamylation patterns and alters microtubule dynamics. **A)** Western blotting of HEK293T cell lysates expressing EYFP-tagged TTLLs or EYFP only shows variable TTLL expression levels. **B)** Western blotting of HEK293T cell lysates expressing EYFP-tagged TTLLs or EYFP only reveals various chain initiation or elongation functions by probing for two tubulin glutamylation epitopes: GT335 for branch point, and B3 for chains longer than two glutamate residues. **C)** Representation of image processing for the analysis of microtubule dynamics via live-cell EB3 imaging. Scale bar = 10 µm. **D-F)** Microtubule dynamics of HEK293T cells co-transfected with EYFP-tagged TTLLs and tdTomato-EB3. Growing microtubule plus-ends were monitored in living cells for 1 min (1 frame per 2s). Graphs show quantifications of microtubule (MT) stability **(D)**, run length **(E)**, and growth rate **(F)**. N = 3, n = 9-11 cells per condition. Shapiro–Wilk test was performed to test for normal distribution of data; statistical analysis was performed by one-way ANOVA with Dunnett’s test for correction of multiple comparisons. ^ns^ non-significance, * P _≤_ 0.05, *** P _≤_ 0.001.

When analyzing the glutamylation patterns, the TTLLs grouped according to their established functional roles. Lysates of cells expressing the initiator TTLL4 showed stronger signal when probed with GT335 antibody, which detects the initial branching point of the glutamate chain. In contrast, lysates of cells expressing the elongators TTLL6 or TTLL11 displayed stronger bands when probed with B3 antibody, which specifically recognizes glutamate side chains with more than two residues. Interestingly, TTLL7 showed both initiation and elongation activities, producing positive signals with both antibodies, whereas TTLL1 showed only barely detectable activity with B3, but not with GT335 antibody (Fig 1B). In sum, transfection of HEK293T cells with our constructs resulted in expression of TTLL enzymes with the expected size and the expected polyglutamylation activity.

### TTLL6 expression decreases microtubule stability

Next, to assess whether expression of individual TTLLs affects microtubule dynamics besides glutamylation, we co-transfected HEK293T cells with EB3-tdTomato and the respective EYFP-TTLL for 2 days, and tracked EB3 comets in yellow fluorescent (i.e. co-transfected) cells (Fig. 1C). Analysis revealed that EYFP-TTLL6, but not any other expressed TTLLs, led to decreased microtubule stability (in terms of duration of traceable comets) compared to cells expressing EYFP alone (Fig. 1D), while microtubule run length remained unchanged (Fig. 1E). Interestingly, this TTLL6-induced microtubule instability was associated with an increase in microtubule growth rate (Fig. 1F). Hence, while TTLL6 had one of the lowest impacts on microtubule glutamylation, it may be an important player for microtubule dynamics.

### LDC10 inhibits TTLL-induced tubulin glutamylation and restores microtubule stability

A TTLL inhibitor identified through a high throughput screening (HTS) campaign at the Lead Discovery Center GmbH was further evaluated as a chemical tool to dissect glutamylation in cells. This hit compound is identified as LDC 10 (unpublished data). To this end, we treated HEK293T cells expressing various EYFP-TTLLs with 10 µM LDC10 for 24 hours and assessed tubulin glutamylation using GT335 and B3 antibodies via Western blotting. Cells treated with LDC10 showed a significant reduction in both monoglutamylation or polyglutamylation compared to vehicle-treated control (Fig. 2A-B). Notably, LDC10 protected microtubules from the destabilizing effects of TTLL6, with LDC10-treated EYFP-TTLL6-expressing cells exhibiting microtubule stability and growth rate levels similar to those in cells expressing EYFP alone (Fig. 2C-E).

**Figure 2.**
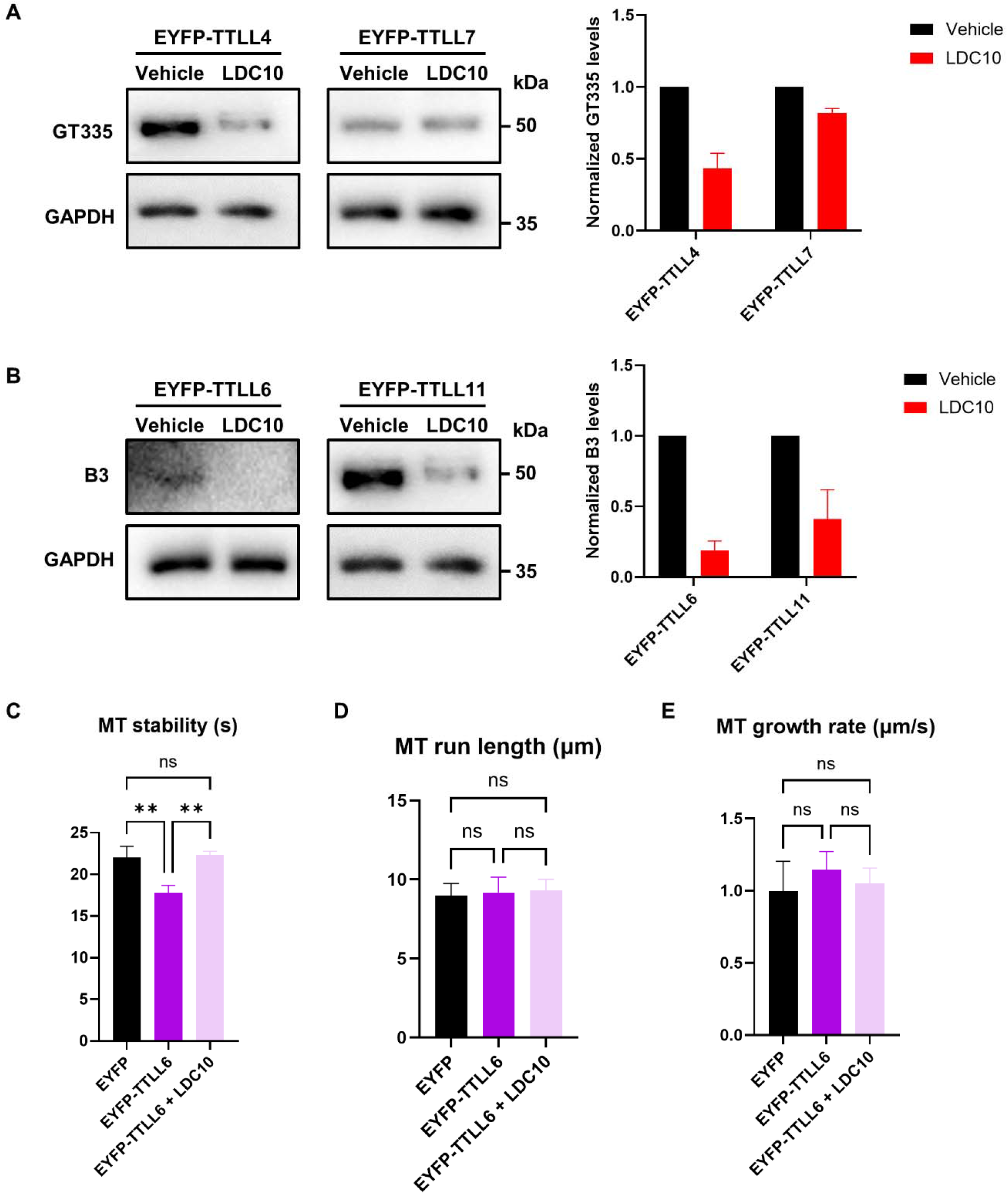
LDC10 inhibits TTLL-induced glutamylation and restores microtubule stability. **A)** Western blotting of HEK293T cell lysates expressing initiators TTLL4 or TTLL7 shows decreased monoglutamylation levels following LDC10 treatment for 24 hours. Images representative of 2-3 Western blots. **B)** Western blotting of HEK293T cell lysates expressing elongators TTLL6 or TTLL11 demonstrates decreased polyglutamylation levels following LDC10 treatment for 24 hours. Images representative of two Western blots. **C-E)** Microtubule dynamics of HEK293T cells co-transfected with EYFP-tagged TTLLs and tdTomato-EB3. Growing microtubule plus-ends were monitored in living cells for 1 min (1 frame per 2s). Graphs show quantification of microtubule (MT) stability **(C)**, run length **(D)**, and growth rate **(E)**. N = 3, n = 9-11 cells per condition. Shapiro–Wilk test was performed to test for normal distribution of data; statistical analysis was performed by one-way ANOVA with Tukey’s test for correction of multiple comparisons. ^ns^ non-significance, ** P _≤_ 0.01.

## Discussion

In this study, we investigated the effects of several human recombinant glutamylating TTLLs by expressing them in HEK293T cells. Compared to EYFP control, EYFP-TTLLs showed low expression levels. Notably, TTLL11 exhibited relatively robust expression and strong polyglutamylation activity, while TTLL4 displayed prominent monoglutamylation activity despite having the lowest expression among all investigated TTLLs. Glutamylation patterns probed at two different epitopes, one marking the branch starting point and the other marking long chains, corresponded with canonical TTLL initiation (TTLL4) and elongation (TTLL6 and TTLL11) functions, with the exception of TTLL7, which showed both activities despite being classified as an initiator. Remarkably, the initiase TTLL4 displayed some elongation activity, and the elongase TTLL11 exhibited minor initiation activity. TTLL1, in contrast, showed barely any detectable polyglutamylation activity, consistent with prior reports that it requires a five-subunit complex to be active (Garnham et al., 2015; Janke et al., 2005).

Glutamylation is known to regulate microtubule stability, dynamics, and function. Therefore, we investigated whether any TTLL would affect microtubule dynamics. Using live-cell imaging of fluorescently tagged EB3 protein, we observed that only TTLL6 negatively impacted microtubule stability while increasing microtubule growth rate. Long glutamate side chains are associated with spastin recruitment and subsequent microtubule destabilization due to microtubule severing (Brill et al., 2016; Lacroix et al., 2010; Roll-Mecak & Vale, 2005, 2008), which could explain decreased microtubule stability after the expression of the elongase TTLL6, and why the initiases TTLL4 and TTLL7 showed no such effect. However, this does not explain the apparent lack of impact on microtubule dynamics by the other elongase TTLL11, especially since it has remarkably higher expression and activity than TTLL6, as seen in our Western blot experiments. One possible explanation is that TTLL6 and TTLL11 modify different sites within the C-terminal tail of tubulin with different spastin recruitment capabilities. Another explanation is that while both TTLL6 and TTLL11 are elongators, the length of the side chain they generate is different. This could be linked to our Western blot results which showed a much stronger long chain band with TTLL11. Glutamate chains containing more than eight residues are known to inhibit spastin severing activity, switching its function to microtubule stabilization instead (Valenstein & Roll-Mecak, 2016). It could be that the strong polyglutamylation induced by TTLL11 in our model stabilizes microtubules, while modest polyglutamylation by TTLL6 is enough to recruit spastin and promote severing and destabilization. A third explanation would revolve around the severing enzyme recruited. Polyglutamylation does not only recruit spastin, but it is also able to recruit katanin, another microtubule severing enzyme (McNally & Vale, 1993; Szczesna et al., 2022). It has been shown before that TTLL6 induced a much stronger katanin activation compared to TTLL11 (Lacroix et al., 2010), suggesting that katanin, and not spastin, could be the primary downstream effector here.

Surprisingly, TTLL6-induced microtubule destabilization was associated with an increased microtubule growth rate. While this may seem contradictory at first, a simple explanation would rely on spastin recruitment. Spastin-mediated microtubule severing has been shown to increase the pool of free tubulin, making more tubulin available for the polymerization of new microtubules, and thus contributing to microtubule regrowth and organization (Aiken & Holzbaur, 2024; Kuo et al., 2019).

Given the therapeutic relevance of glutamylating TTLLs in cancer and neurodegenerative diseases (Das et al., 2014; Rogowski et al., 2021; Wu et al., 2022), finding new TTLL inhibitors is critical. We tested LDC10, a novel TTLL inhibitor, for its efficacy in reducing TTLL-induced glutamylation and reversing TTLL6-mediated microtubule instability. Our results show that LDC10 is able to inhibit TTLL4-induced monoglutamylation and, to a lower extent, TTLL7-induced monoglutamylation, as well as polyglutamylation generated by TTLL6 and TTLL11. Notably, LDC10 completely reversed the microtubule destabilizing effect of TTLL6, restoring stability and growth rate to control values. Although LDC10 shows initial promise in these microtubule stability assays as a tool compound, together with the remaining hits from the TTLL4 HTS, it is still under medicinal chemistry-based optimization to enhance specific potency against inhibition of TTLL4 and to improve its lead-and drug-likeness.

In conclusion, we showed direct detrimental effect of human TTLL6 on microtubule dynamics in HEK293T cells and presented LCD10 as a chemical tool compound that inhibits glutamylation and is capable of mitigating the associated negative effects.

## Availability of data and materials

The datasets of the current study are available from the corresponding author on reasonable request.

## Authors’ contributions

Study design: MAAK, HZ. Experimental work, data acquisition and analysis, and manuscript writing: MAAK. Drug development: PJ, KKl. All authors proofread, commented on, and approved the final manuscript.

## Acknowledgments

We thank Daniel Adam and Jennifer Klimek (CMMC and Institute of Human Genetics, University Hospital Cologne, Cologne, Germany), and Tamara Wied (current address: Max Planck Institute for Biology of Ageing, Cologne, Germany) for the technical help and stimulating discussions. We thank Carsten Janke (Institut Curie, Paris, France) for the provision of the plasmids.

